# Fast and reliable ancestral reconstruction on ancient genotype data with non-negative Least square and Principal Component Analysis

**DOI:** 10.1101/2024.05.06.592724

**Authors:** Luciana de Gennaro, Ludovica Molinaro, Alessandro Raveane, Federica Santonastaso, Sandro Sublimi Saponetti, Michela Carlotta Massi, Luca Pagani, Mait Metspalu, Garrett Hellenthal, Toomas Kivisild, Mario Ventura, Francesco Montinaro

## Abstract

The history of human populations has been strongly shaped by admixture events, contributing to the patterns of observed genetic diversity across populations. Given its significance for evolutionary and medical studies, many algorithms focusing on the inference of the genetic composition of admixed populations have been developed. In particular, the recent development of new ancestry estimation methods that consider the fragmentary nature of ancient genotype data, such as the f-statistics family and its derivations, have radically changed our understanding of the past. F-statistics capture similar genetic similarity information as Principal Component Analysis (PCA), which is widely used in population genetics to quantify genetic affinity between populations or individuals. In this study, we introduce ASAP (ASsessing ancestry proportions through Principal component Analysis) method that leverages PCA and Non-Negative Least Square (NNLS) to assess the ancestral compositions of admixed individuals given a large set of populations. We tested ASAP on different simulated models, incorporating high levels of missingness. Our results show its ability to reliably estimate ancestry across numerous scenarios, even those with a significant proportion of missing genotypes, in a fraction of the time required when using other tools. When harnessed on Eurasia’s genotype data, ASAP helped replicate and extend findings from previous studies proving to be a fast, efficient, and straightforward new ancestry estimation tool.

## Introduction

The history of human populations has been strongly shaped by past admixture events that cumulatively have contributed to patterns of genetic diversity observed today (Hellenthal et al., 2014; Montinaro et al., 2015). Several multidisciplinary studies proved that virtually all human populations have interacted throughout their history in complex demographic scenarios, including migration and admixture (Busby et al., 2016; Ongaro et al., 2019; Patterson et al., 2012; Schlebusch et al., 2012). These interactions resulted in a sudden or gradual transfer of genetic material, generating new groups different from their sources (Hellenthal et al., 2014). Given its significance for evolutionary and medical studies, many algorithms focusing on the inference of the genetic composition of admixed populations have been developed. In this context, it has been shown that using phased genotype data can offer higher-resolution description of genetic population structure (Haak et al., 2015; Hellenthal et al., 2014; Lazaridis et al., 2016; Leslie et al., 2015; Montinaro et al., 2015; Narasimhan et al., 2019; Ongaro et al., 2019; Pankratov et al., 2020; Skoglund et al., 2017).

However, existing methods often present limitations when dealing with low-coverage ancient DNA (aDNA) data. Algorithms using haploid-called genotypes to estimate allele frequencies and allele sharing probabilities at limited numbers of overlapping variant positions have been designed to meet these challenges.

Among the others, qpAdm (Haak et al., 2015; Harney et al., 2021) is one of the most widely used approaches on ancient data (Damgaard et al., 2018; Haber et al., 2017; Hajdinjak et al., 2018; Harney et al., 2018; Lazaridis et al., 2017, 2016; Narasimhan et al., 2019; Olalde et al., 2018; Skoglund et al., 2017), given its ability to deal with the challenges of aDNA data and model admixture events involving multiple sources (Haak et al., 2015; Harney et al., 2021; Patterson et al., 2012). This tool takes advantage of the fact that genetic variation within a specific population can be summarised by comparing its allele frequencies to those of three additional groups using a “treeness test” belonging to the F-statistics family, the *f4* metric (Harney et al., 2021; Patterson et al., 2006).

For a given target population T, a set of putative sources of admixture P_i_, and a set of “right populations” R_i_ with different relationships to P_i_, qpAdm builds a matrix A of *f4* in the form (T, X, R_1_, R_i_), in which X can alternatively be T or a P_i_ population. Given that any *f4* in the state (T, T, R_1_, R_i_) is 0, qpAdm solves the equation *w.A=0,* where w are the admixture coefficients (weights), assuming that their sum is equal to 1 (Haak et al., 2015).

QpAdm framework can be iterated multiple times to test several scenarios, allowing the evaluation of the models based on their p-values. However, sifting through all possible proxy sources and the right populations for an admixture event can be overwhelming. In addition, a recent survey has shown that, depending on the approach and the quality of the genetic data analyzed, qpAdm may suffer from high false discovery rates, adding substantial uncertainty to the interpretation of the results of admixture inference (Eren Yüncü et al., 2023).

A similar approach, introduced by Haak et al. 2015, but less frequently employed, uses a Non-Negative Least Square (NNLS) approach on a matrix of *f4*s in the form *f4*(X, R_1_, R_i_, R_j_), where X is either T or any P population (Haak et al., 2015; Lazaridis et al., 2016).

*F*-statistics results broadly recapitulate genetic relationships emerging from Principal Component Analysis (PCA) (Peter, 2022), widely used in population genetics to quantify genetic affinity between populations or individuals, including ancient ones.

There is indeed a geometric relationship between the two metrics, although they are based on different statistical principles: the f-statistic is based on the measurement of the branch lengths of a hypothetical tree in which the analyzed populations are related, while PCA reduces the dimensionality of the data while maintaining the maximum variance present among individuals. In detail, considering four populations A, B, C, and D projected in a PC space, the *f2*(A, B) is correlated with the Euclidean distance between A and B computed in PC coordinates, while the f3(A; B, C) will be proportional to the orthogonal projection of A-B on A-C. Similarly, the *f4*(A, B; C, D) will be related to the orthogonal projection of A-B onto C-D (Peter, 2022). Moorjani et al. 2011 showed that f4 ratios can be used to estimate the rate of admixture (Moorjani et al., 2011).

Considering these results, it is, in principle, possible to use PC coordinates to infer admixture proportions of a target population using a set of sources. Different attempts and approaches have recently been proposed using principal components (Conley et al., 2023).

In this study, we present ASAP (ASsessing Ancestry proportions through Principal component analysis), in which we aim to leverage PCA and NNLS to assess the ancestral compositions of admixed individuals given a large set of populations. We test ASAP on different simulated models, incorporating high levels of missingness. We show its ability to reliably estimate ancestry across numerous scenarios, even those with a significant proportion of missing genotypes, in a fraction of the time required when using other tools.

## RESULTS

### ASAP workflow and datasets

Here we provide an overview of the methodology implemented in ASAP (Fig. 1). We simulated a set of 20 unadmixed and 16 admixed populations (Supp. Table 1). For each admixed group, we simulated an admixture event involving two or three sources (Molinaro et al., 2021) with minor source contribution ranging from 5 to 40%, to test ASAP performance in various conditions and settings, accommodating a wide range of routinely performed approaches. For each of the true sources, we also simulated a sister group that split 3 thousand years ago (KYA) to mimic a proxy source: a group related to the real admixing source but not the direct contributor to the admixture event. These proxy populations allowed us to test whether ASAP could infer the closest proxy sources to the admixing populations.

**Fig. 1.**
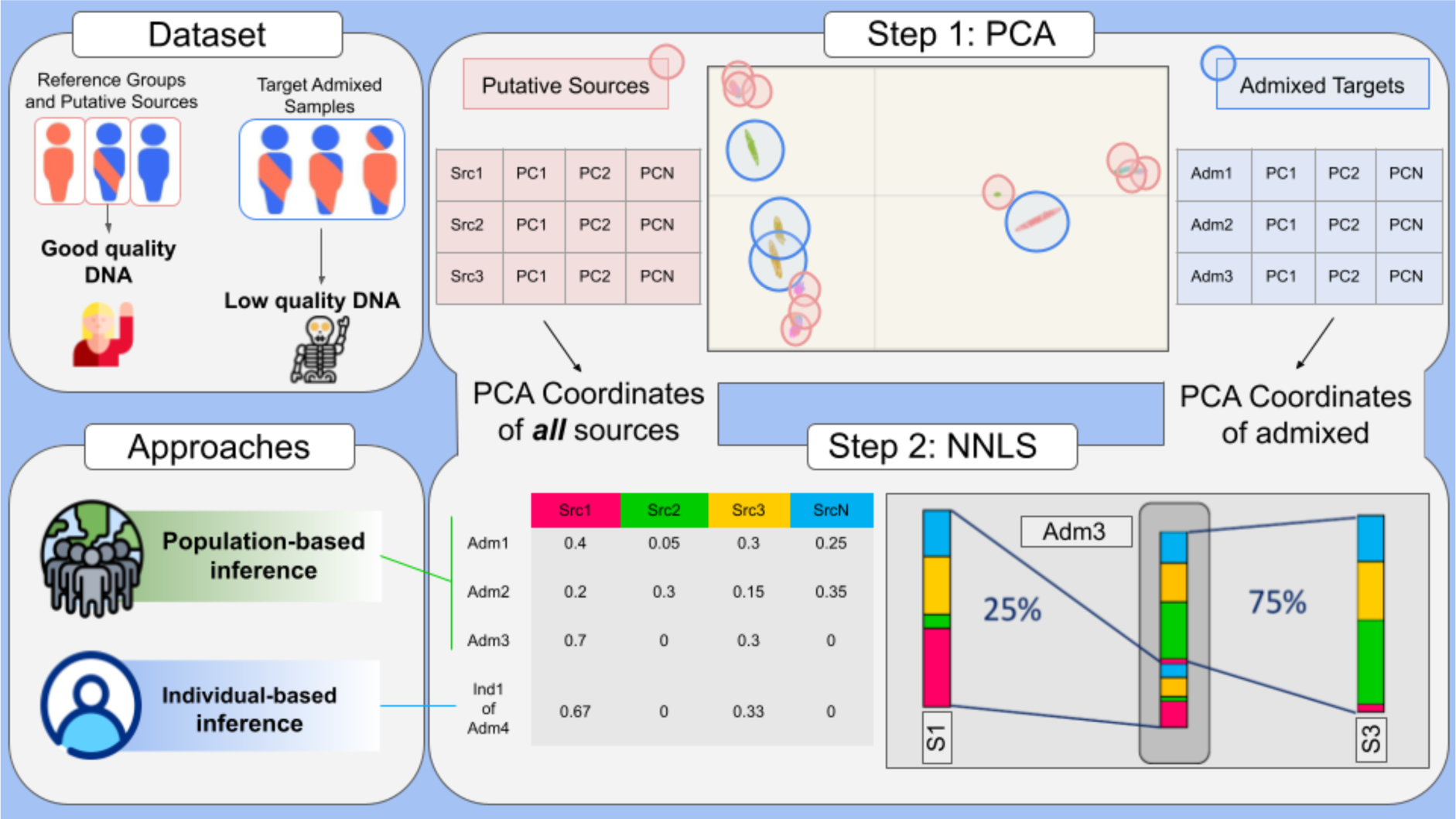
Schematic representation of ASAP workflow. ASAP harnesses Non Negative Least Square using individual or population Principal Component vectors. PCA analysis can be carried out on no-missing genotype data, or using different approaches which accommodate different degrees of missingness.

Specifically, we simulated admixture events between groups with different degrees of affinity, from highly divergent to closely related populations, with pairwise Fst between populations ranging from 0.01 to 0.23, including bottleneck events, expecting a lower assignment accuracy in cases where the source groups are genetically closer (Molinaro et al., 2021). For each scenario, we tested our approach on the average Principal Component (PC) coordinates from each admixed group (population-wise approach) and on each admixed individual separately (individual-wise approach).

We initially tested ASAP performance considering as putative admixture sources the entire panel of the true sources or their sister groups (‘proxy sources’), even though only two (two-way admixture) or three (three-way admixture) sources were used to simulate the admixture event. We then ran PCA where the PC space was built by the true sources and their respective sister groups on the first 10 components, while all admixed groups were projected onto it. The number of components was selected after running a preliminary assessment of ASAP performance as described in Supplementary Text 1. Subsequently, using NNLS, we modeled the average PC coordinates across individuals of each admixed group as a mixture of those of all the available sources, considering as sources either the true or the sister groups panel. Standard Errors (SE) were estimated using a chromosome-based jackknife approach (Busing et al., 1999; Montinaro et al., 2015), as described in the methods section.

### Data and source availability

Our approach to ASsess Ancestral composition using Principal Component Analysis and NNLS (ASAP) is available as an R package at https://github.com/lm-ut/ASAP.

### Results on Simulated genotypes with no missingness

#### ASAP performance with true sources

In the population-based approach, for all the 16 simulated admixed populations, ASAP successfully assigned the main ancestry components to the true sources that contributed to the admixture event despite the large panel of potential source groups available. The ancestry proportions of the true source groups (Supp. Table 2a) yielded a maximum error of 0.014 and a maximum jackknife standard error (SE) of 0.012, when two sources contributed to the target population (Fig. 2A-C). Minor additional contributions were assigned to other groups but never exceeding 0.004 (Pop 8 in Fig. 2A). In these cases, the additional ancestral component was assigned to groups closely related to the true source (Fst ≤ 0.01, (Molinaro et al., 2021)). In three-way admixed populations (Pop 15 and Pop 16 in Fig. 2D-E), the true sources are always recognized with a maximum error of 0.010 (SE < 0.01).

**Fig. 2.**
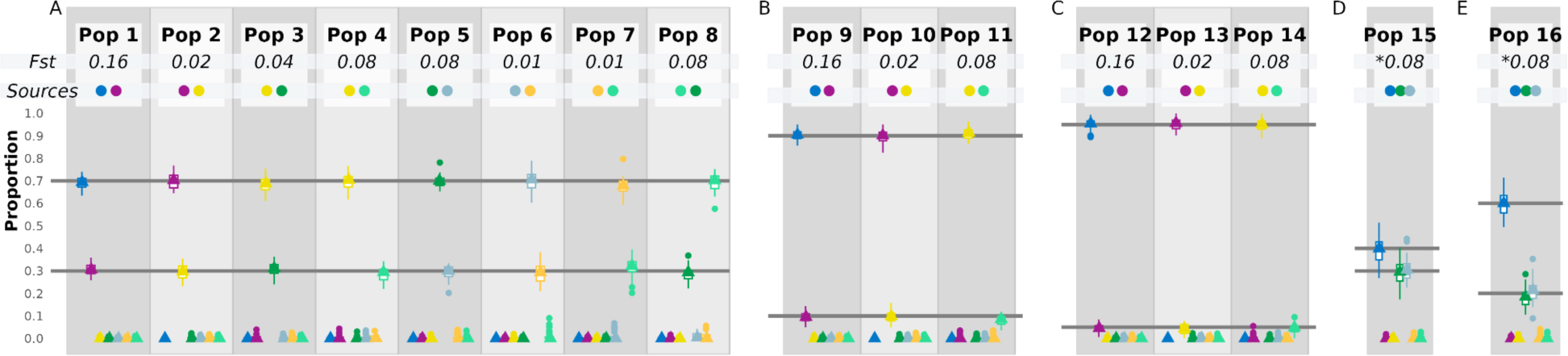
ASAP assignment using true sources for each admixed group; triangles and boxplots show the population and individual ancestry estimation, respectively. On the upper part of the panel the *Fst* values (minimum values are marked with *) and the *Sources* that contribute to the admixture event in the simulated populations are shown: A) populations obtained from the combination of two sources with proportions 70-30%; B) admixed populations obtained from the combination of two sources with proportions 90-10% and **C)** 95-5%. **D)** Three-way admixed populations generated by combining three sources with 40-30-30% and **E)** 60-20-20% proportions.

The individual ancestry assignment estimation for (Supp. Table 2) 70-30% admixed populations (Pop 1-8) shows an average error lower than 0.029 when the admixing sources split more than 9 KYA (Kilo Years Ago), which increases (0.038) when the simulated population split was less than or equal to 9 KYA (Pop 6 and 7).

Admixed populations with lower source contributions (Pop 9-14) record an average error of a maximum of 0.022. In this case, lower error values are observed for populations with a less recent split (for example Pop 9 and Pop 12). The highest average error in the individuals-based analysis is observed in the three-way admixed populations (Supp. Table 2b). An over/underestimation exceeding 0.05 of the assigned contribution to the main sources in the 32% of individuals is recorded. Only one individual has an error larger than 0.1.

#### ASAP performance with proxy sources

We evaluated ASAP performance on a PCA where the admixed groups were projected onto the PC space built on all the remaining populations. We then modeled the admixed groups as a mixture of all the proxy sources only. As we knew which simulated proxy group was indeed the sister group of the real admixing source, we calculated the assignment error by considering differences in observed and expected ancestry proportions and whether ASAP could indeed select the closest sister group. In this scenario, for all the 16 tested populations the proxies of the real sources were recognized without ever assigning even minimal contributions to other populations (Supp. Table 2).

For two-way admixed populations with proportions of 70-30%, the average error is 0.033 (SE < 0.009, Supp. Table 2C). Generally, the error estimates tend to be larger when the admixing source populations (Pop 3, 4, 6, 7 in Fig. 3) are characterized by a higher genetic similarity due to recent split times and bottleneck events. However, the error never exceeds 0.057 (Pop 4).

**Fig. 3.**
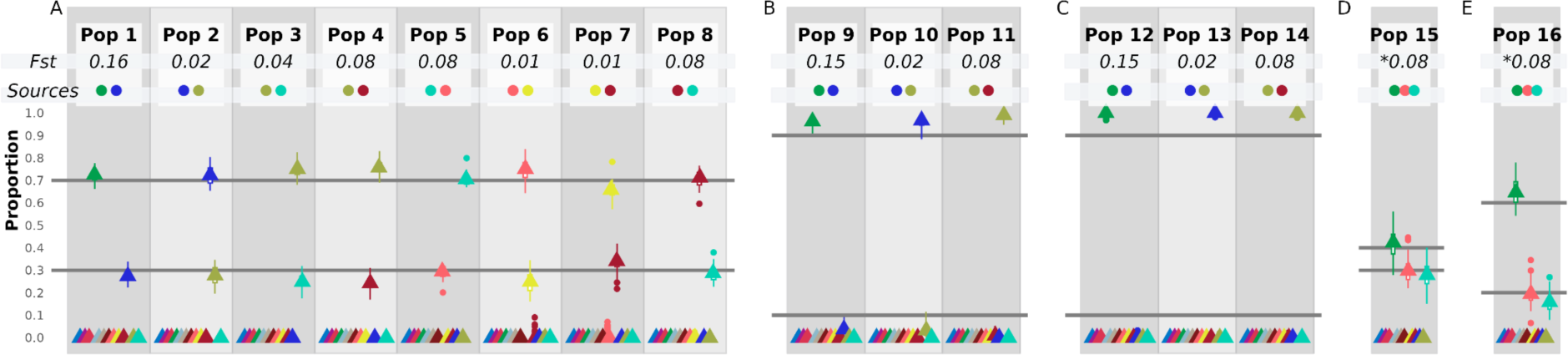
ASAP assignment using proxy sources for each admixed group. Triangles and boxplots show the population and individual ancestry estimation, respectively. On the upper part of the panel the *Fst* values (minimum values are marked with *) and the proxy *Sources* that contribute to the admixture event in the simulated populations are shown: A) populations obtained from the combination of two sources with proportions 70-30%; B) admixed populations obtained from the combination of two sources with proportions 90-10% and C) 95-5%. D) Three-admixed populations generated by combining three sources with 40-30-30% and E) 60-20-20% proportions.

The ASAP accuracy is also robust in the case of three-way admixed populations, with a maximum error of 0.031 (SE < 0.0087).

The overestimation of the major component becomes more important in strongly imbalanced contribution cases. When the contribution of the minor source is 10% (Fig. 3B), the minor contribution is underestimated (Pop 10-11 in Fig. 3B). For populations where the minor source contributed 5%, ASAP completely misses the minor source contribution and assigns the total of the ancestral component to the main source (SE < 0.007) (Fig. 3C).

In individual-based inferences (Supp. Table 2D), ASAP correctly assigns ancestral proportions in the two-70%-30% admixed individuals. The estimates obtained for the 50 individuals within each group are characterized by a maximum average error of 0.0582 (Pop 6, Fig. 3A).

For admixed populations with minor source contributions of 10% and 5%, the contribution of the minor sources is underestimated or completely missed. In detail, for populations 9 and 10, the average estimated minor contribution is 0.038 and 0.034, with 29% of individuals showing less than 2%. On the other hand, in population 11, the average minor contribution is 0.016, with 64% of individuals showing less than 2%. No relevant contribution from other sources is recorded. For populations 12, 13 and 14, 85% of individuals are modeled as unadmixed, with the remaining individuals showing an average minor contribution of 1.3%.

For the three-way admixed populations, ASAP is always able to recognize the correct sources and assigns them the right proportions with a maximum error of 0.05 (Pop 16, Fig. 3E) due to the fact that for some individuals there is a slight overestimation of the main source (AFR) at the expense of the Asian source, one of the other two minor sources.

### ASAP performance using pseudo-ancient data

We tested ASAP using pseudo-haploid samples, simulated by introducing different degrees of missing genotypes (up to 50%) and pseudo-haplodised (see materials and methods) mirroring the fragmentary nature of data commonly adopted in ancient DNA studies. We tested ASAP on a PCA where the PC space is built by the diploid genomes of the proxy sources, onto which we projected the pseudo-haploid genomes of the admixed groups and all possible true sources. In this scenario, we tested whether ASAP could model the admixed groups with the pseudo-haploid true sources.

ASAP correctly detects the closest admixture sources even in a large panel of putative donors, despite the target and source samples being pseudo-haploid and containing missing genotypes (Supp. Table 3, Fig. 4). Indeed, the average maximum assignment error is 0.033. In this case, ASAP always identifies the true sources and assigns a marginal additional component to other sources (maximum 0.004). Furthermore, the jackknife SE is also generally low, with a maximum of 0.017 (Supp. Table 3) seen in the admixed population whose sources split more recently (7.5 KYA). Even when single samples are targeted, the true sources are generally recognized and the major source ancestry assignments show an average error of 0.039. Despite the low average error, the maximum per sample error can reach 0.248, caused by the misassignment to the most closely related group to the sources (Fst % 0.01,(Molinaro et al., 2021)).

**Fig. 4.**
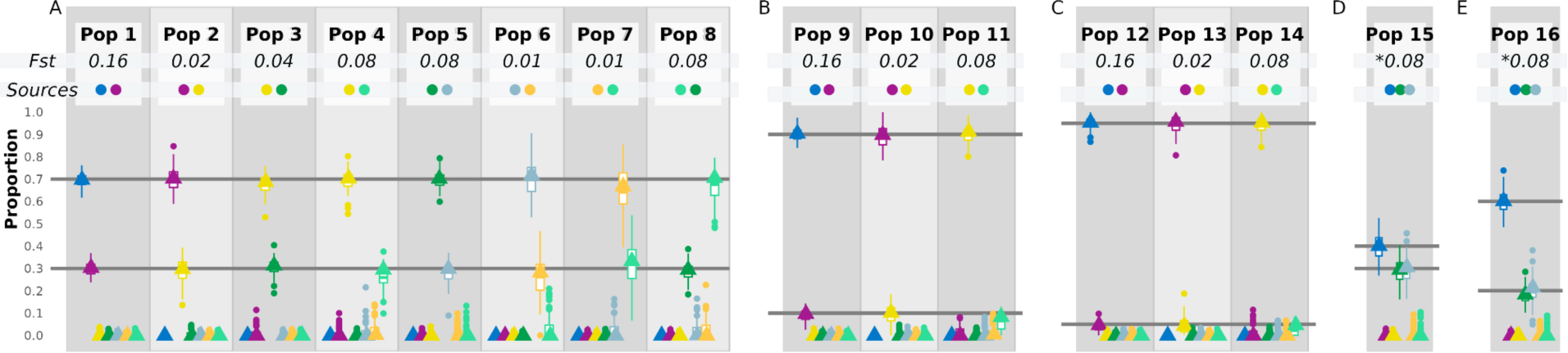
ASAP assignment using pseudo-haploid simulated data and modeling each admixed group as a mixture of all the available proxy sources. Triangles and boxplots show the population and individual ancestry estimation, respectively. n the upper part of the panel the *Fst* values (minimum values are marked with *) and the proxy *Sources* that contribute to the admixture event in the simulated populations are shown: A) populations obtained from the combination of two sources with proportions 70-30%; B) admixed populations obtained from the combination of two sources with proportions 90-10% and C) 95-5%. D) Three-admixed populations generated by combining three sources with 40-30-30% and E) 60-20-20% proportions.

#### ASAP performance with limited reference genetic variation availability

We tested ASAP in a scenario where only the proxy, but not the true aDNA-like pseudo-haploid sources of the admixture, were available. The rationale behind this analysis is to mimic the lack of true mixing sources when exploring ancient DNA datasets while leveraging the availability of diploid genomes to build the PC space. In this case, an initial hypothesis of the demographic history of the admixed group is required, given that we are subsetting the donor panel to two or three putative source groups.

In this test, we projected onto the PC space the pseudo-haploid genomes of i) the target admixed group and ii) the closest proxy sources of each real source, two proxy sources in case of a two-way admixture, and three for the three-way admixture. The PC space was built with the diploid genomes of the remaining proxy sources. We modeled the target admixed group’s relative admixture proportions given the projected proxy sources, relying on a limited donor panel of two or three groups.

Given the large error in individual analysis, mostly due to the lack of a proper reference dataset, we focused on the population-based approach (Fig. 5). In such a scenario (complete results available in Supp. Table 4), error estimates are lower than 0.043 for all groups whose sources diverged more than 24 KYA (Pop 1, 4, 5, 8, 9, 11, 12, 14). For the only group whose sources split 24 KYA (Pop 3), the error increases to 0.11. In contrast, for all the other groups with closer sources, the error estimates range between 0.16 and 0.58, with jackknife SE estimation following the same pattern (Supp. Table 4).

**Fig. 5.**
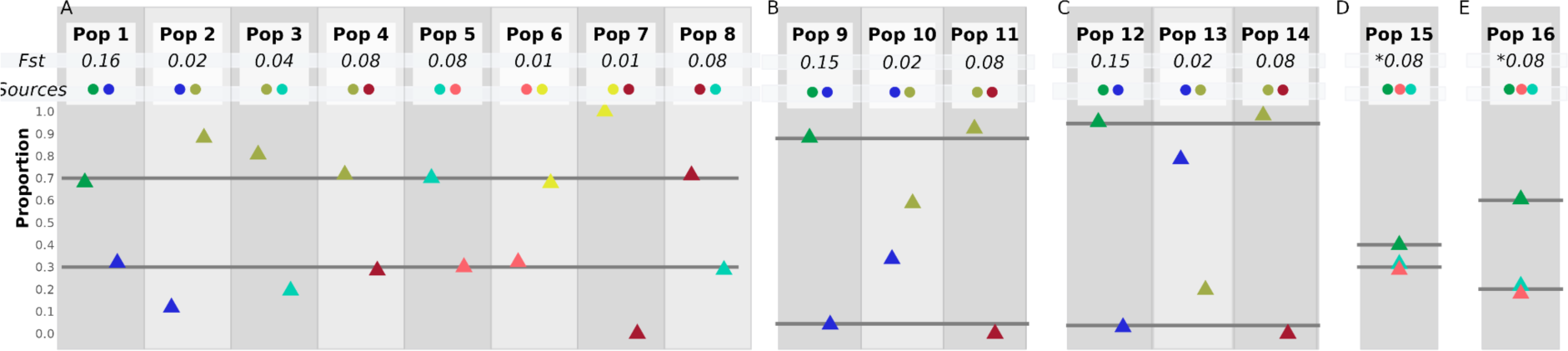
ASAP performance with limited reference genetic variation availability. Only population-based inferences are shown.n the upper part of the panel the *Fst* values (minimum values are marked with *) and the proxy *Sources* that contribute to the admixture event in the simulated populations are shown: A) populations obtained from the combination of two sources with proportions 70-30%; B) admixed populations obtained from the combination of two sources with proportions 90-10% and C) 95-5%. D) Three-admixed populations generated by combining three sources with 40-30-30% and E) 60-20-20% proportions.

### Benchmarking ASAP versus existing global ancestry inference tools

We compared ASAP with qpAdm, Rye, and Unlinked-ChromoPainter NNLS, which harness f4-statistics, PCA, and a modified Li and Stephens model with infinite recombination between SNPs for the ancestry composition inference, respectively (Conley et al., 2023; Haak et al., 2015; Harney et al., 2021; Li and Stephens, 2003). We compared the accuracy in estimating the ancestral proportions of the four approaches using the pseudo-haploid genomes of both the target admixed samples and the true sources of the admixture. Our method behaves similarly to the others (Fig. 6A-C); the correlation of ancestry assignments (Fig. 6D) of ASAP, qpAdm, and Rye is higher than 0.95 (ASAP vs qpAdm R2 = 0.968, p-value < 10e-6; ASAP vs Rye R2 = 0.998, p-value < 10e-6, ASAP vs CP R2 = 0.985). Among the four harnessed algorithms, qpAdm is characterized by the highest average error, and all four approaches show a lower accuracy for the admixed populations characterized by a subcontinental admixture, in which the two admixing sources are generically close (Fst ≤ 0.01) (Molinaro et al., 2021; See Suppl. Table 5).

**Fig. 6.**
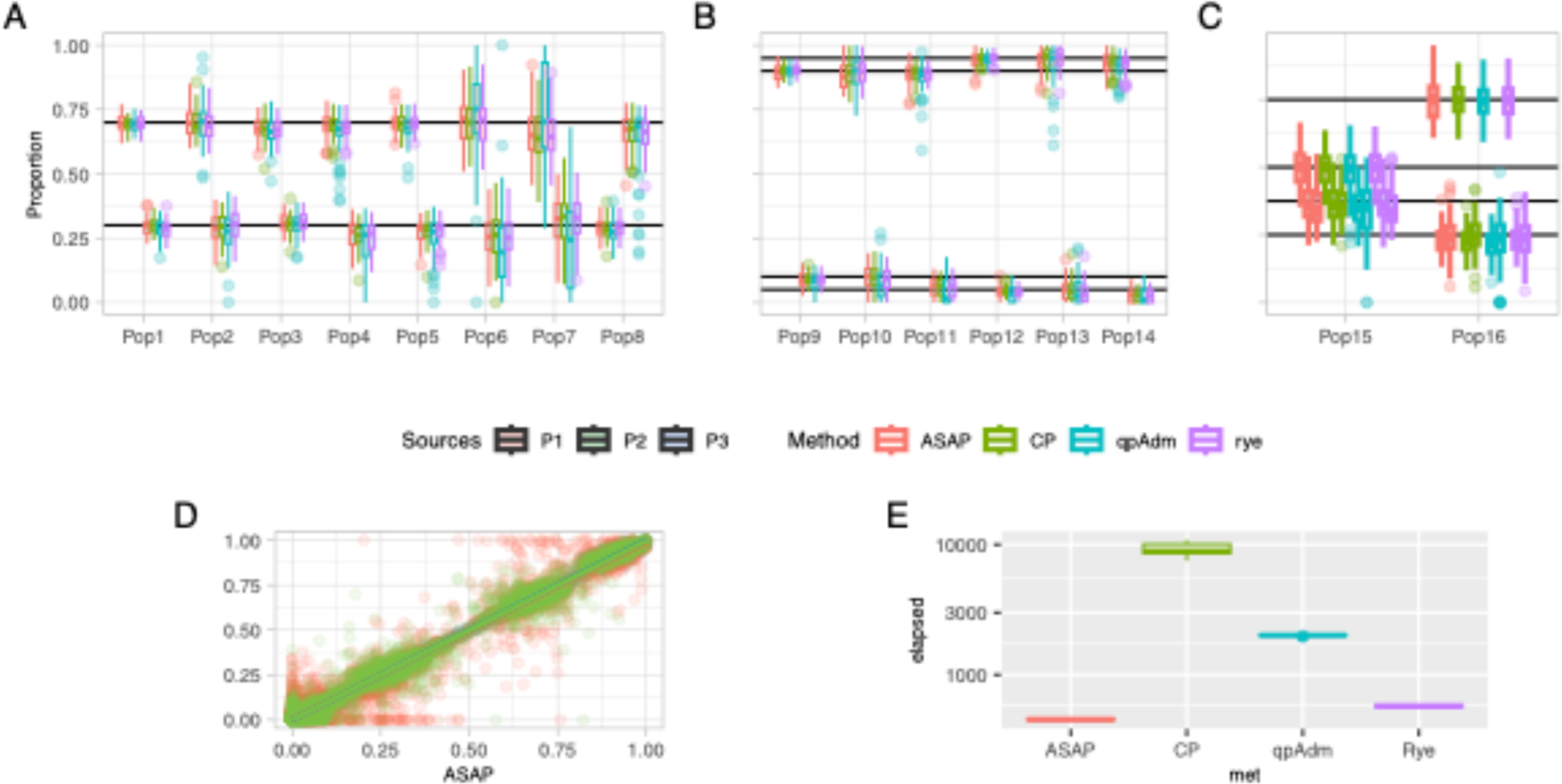
Comparison between ASAP (red), ChromoPainter NNLS (CP, green), qpAdm (blue), and Rye (purple) modeling the ancestral proportions of pseudo-haploid admixed populations given a set of pseudo-haploid sources; **A)** ancestry proportions for simulated populations with 70%-30% sources’ contribution; **B)** ancestry proportions for simulated populations with 90% - 10% (Pop 9-11) and 95%-5% (Pop 12-14) sources’ contribution; **C)** ancestry proportions for three-way admixture simulated populations with 40%-30%-30% (Pop 15) and 60%-20%-20% (Pop 16) sources’ contribution; **D)** correlation of the ancestry proportion assignment between CP, Rye and qpAdm in the y-axis and ASAP on the x-axis and **E)** computational time for a subset of 100 individuals.

We also compared the computational speed of each framework (Fig. 6E) replicating (10 iterations) the ancestry inference for the same set of 100 individuals. For ASAP and Rye, we included the PC (10 analysis) time, while for qpAdm we took into consideration the estimation of the f2. ASAP outperforms the other methods: ASAP computational time stands at 454 seconds (s) (sd = 5 s), while Rye reaches 575 s (sd = 14.1 s), qpAdm 2,011 s (sd = 22.399 s), and ChromoPainter 156 minutes and 23 seconds (9,383 s, sd = 1148 s).

### ASAP performance on real data

We tested ASAP on real data using a dataset of different ancient Eurasian populations (Lazaridis et al., 2022). We projected 1,380 ancient individuals into the first 10 Principal components inferred using 1,668 present-day individuals. Following Lazaridis et al. 2022, we applied ASAP on 1,350 target individuals, using five putative sources: Western Hunter-Gatherers (WHG), Caucasus Hunter-Gatherers (CHG), Eastern Hunter-Gatherers (EHG), Anatolia Neolithic and Levant Neolithic.

The ancestry compositions captured by ASAP on real data show a significant correlation (R=0.92, p<.0001) with F4admix results obtained in the original paper (Lazaridis et al., 2022), confirming the reliability of ASAP in real-world scenarios (Fig. 7, Suppl. Table 6).

Moreover, we explored the individual ancestral composition of specific geographic locations in different time transects as in Lazaridis (Lazaridis et al., 2022). This enabled us to pinpoint the emergence of ancestral influences across different geographical regions and prehistoric periods. First, we examined the Anatolian region and confirmed an increase in Caucasus/Levantine ancestries around 3,000 BCE, accompanied by a subsequent reduction in local Anatolian ancestry (Fig. S1). Then, we confirmed the introduction of CHG-related ancestry into Steppe populations around 5,000 BCE, alongside the absence of Anatolian ancestry in this region prior to 3,000 BCE. We did not observe an increase in Levantine PPN ancestry, suggesting that most Eastern influence is associated with Anatolia Neolithic ancestry. Our approach corroborates again the complex genetic composition observed within the Yamnaya cluster, characterized by consistent CHG admixture (Fig. S2).

ASAP analysis further identified a less pronounced overrepresentation of CHG ancestry if compared to EHG ancestry in Aegean Bronze Age populations. This observation suggests significant gene flow occurring after the Neolithic period, particularly during the Early Bronze Age, across the Aegean and Balkan Peninsula regions (Fernandes et al., 2020; Raveane et al., 2022, 2019; Saupe et al., 2021) (Fig. S3). Similar trends were also observed in Italy, where Iron Age Southern Italian samples exhibited the highest frequency of Caucasus hunter-gatherer ancestry, found almost absent in Central Italian Etruscans Fig. S4 (Aneli et al., 2022).

**Fig. 7.**
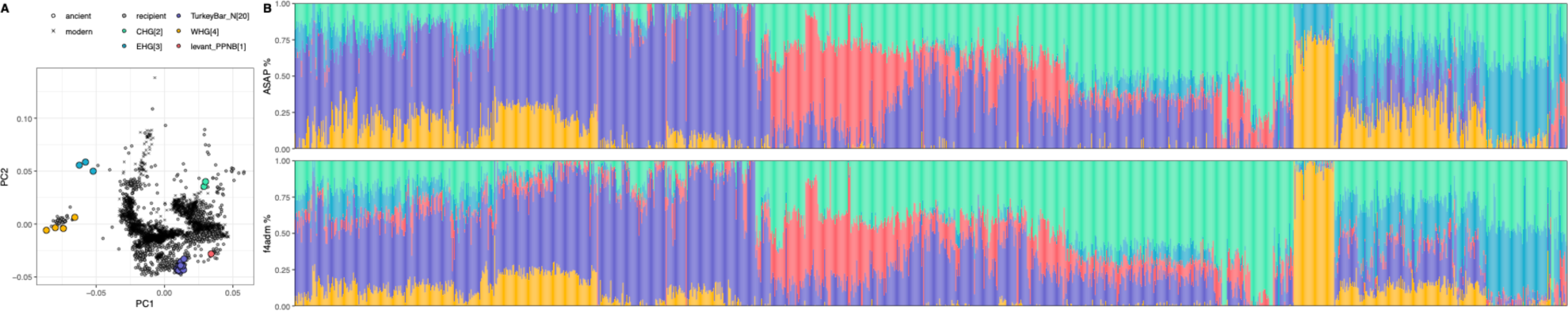
Ancestry inference using ASAP on the aDNA dataset; **A)** PCA used as input by ASAP. **B)** Admixture plot displaying ancestry proportions for 1,350 ancient individuals (x-axis ordered by k-mean cluster numbers computed on ASAP inferred proportions): the upper panel has been estimated using ASAP, while the lower panel shows estimates reported in the original publication using F4admix (Lazaridis et al., 2022).

Although overall there is a high correlation between the two inferences, we observed 273 (out of 6,750) highly discordant estimates (HDE), in which the ancestral proportion difference exceeds 0.2.

1. When considering Western Hunter-Gatherer ancestry, we observed a correlation Pearson correlation coefficient R=0.9 (p-value<1e^-4^) and 53 HDEs. Many of them include Hunter-Gatherers from Serbia and Romania, which are modeled by ASAP as approximately 70% WHG with the remaining ancestry mainly assigned to EHG, while 90% WHG and 10% EHG were estimated by Lazaridis et al. 2022 (Lazaridis et al., 2022). These samples were first published by Mathieson et al. 2018, who described them as a combination of WHG and EHG using qpAdm (although the estimates are associated with very low p-values) and D-statistics (Mathieson et al., 2018).
2. Concerning the ancestry of Turkey Barcin Neolithic individuals, commonly known as Anatolian Neolithic (AN) (Lazaridis et al., 2022), we observed a correlation R=0.95 (p < 1e-4) and 59 HDEs. Nevertheless, a few individuals exhibit a substantial discrepancy in AN ancestry proportion between the two compared methods, making it challenging to determine which of the two approaches has the highest performance. For example, ASAP estimates higher Anatolian Neolithic ancestry for some Mycenaean individuals (Lazaridis et al., 2017) while F4admix gives higher Anatolian ancestry for an Iron Age individual from Lebanon (Haber et al., 2020) (Suppl. Table 6). Both methods can be inaccurate in some cases, as shown by comparisons with previous studies.
3. For Iran Neolithic/CHG ancestry, we observed a correlation R of 0.95 (p-value<1e-4) and 47 HDEs. Most (20) are related to populations from Chalcolithic and Bronze/Iron Age Near East (Iran and Lebanon) individuals. For example, seven Bronze Age individuals from Shahr I Sokhta are modeled as having a substantially smaller Iran Neolithic/CHG ancestry for ASAP estimations (mean=0.65) compared to F4admix (mean=0.94). In Narasimhan et al. (Narasimhan et al., 2019), when Shahr I Sokhta individuals are modeled using qpAdm, they show on average 0.66 IN/CHG (SD=0.05).
4. In the case of EHG, we noted 40 HDEs and an R value of 0.86 (p-value<1e-4). As for WHG, most of the HDEs are related to Hunter-Gatherers from the Iron Gates regions of Serbia, Romania, for which ASAP estimates a higher proportion of EHG when compared to Lazaridis et al. (Lazaridis et al., 2022). Furthermore, in four Bell beaker individuals from Germany, France, and England, ASAP estimates a very low proportion of such ancestry.
5. We observed 74 HDEs and a correlation R of 0.88 (p-value<1e-4) for Levant Neolithic ancestry. Most of the HDEs are related to ancient individuals from the Near East, for which estimates of Levant PPN are always higher than those inferred by Lazaridis et al. (Lazaridis et al., 2022). These results align with previous estimates on the same samples. For example, for the individual I3832, which was modeled as 0.58 Levant PPN and 0.42 Iran Chalcolithic using LINADMIX in its original publication (Agranat-Tamir et al., 2020), ASAP estimated the Levant PPN proportion at 0.77, which was 0.38 when using F4admix (Lazaridis et al., 2022). A possible explanation for this discrepancy is related to the fact that in (Lazaridis et al., 2022), the same individual is modeled to have approximately ∼0.2 related to AN. Similarly, ASAP’s ancestral composition for individuals from Roman and Iron Age from Lebanon are in line with previous DyStruct inferences (Haber et al., 2020). Furthermore, F4admix (Lazaridis et al., 2017) estimated a substantial proportion of Levant PPN ancestry in two Greek and one Italian Bronze Age samples, in contrast with a series of findings on the same or similar individuals. All these samples are characterized by a missingness rate higher than 40% (I9006), suggesting that in some cases ASAP might be less biased than F4admix estimates.

We also used ASAP to test the robustness of the support behind the hypothesis that the WHG contributions of British farmers came mostly from continental WHG rather than local British WHG (Brace et al., 2019). Therefore, modeled different European Neolithic samples as a mixture of WHG and AN, as in Brace et al., 2019 (Brace et al., 2019). We confirmed that WHG proportions in Iberian Early Neolithic samples similar to those in British Neolithic samples, suggesting a common WHG source (Supp. Table 7). We then tested models with pairwise WHG individuals as possible sources and a single Anatolian Neolithic population (see method). We found that both Iberian and British samples consistently preferred Bichon or Villabruna-associated samples as WHG source, indicating their close relationship to the true source and confirming a shared origin (Supp. Table 8) (Fu et al., 2016).

## Discussion

We present ASAP, a global ancestry exploration approach based on PCA and NNLS, that allows an accurate estimation of ancestry proportions in admixed groups or single samples. The approach leverages how the location of samples on the PC space can be related to the mean time of coalescence between pairs of samples (McVean, 2009), and to the recent observation that PC vectors are strongly related to *f4* metrics (Peter, 2022). Specifically, in the case of an admixture event, samples will fall along a gradient and their putative admixture sources will be placed at the ends of the gradient (McVean, 2009; Patterson et al., 2006). Our approach exploits the relative coordinates of the admixed samples and the ones of the putative sources in the PC space and summarises the ancestry proportions of the targets through NNLS.

An advantage of the method is that it can leverage the entire PC space, allowing a large panel of donors to model a given target admixed group or multiple parallel analyses in case several different target groups and their relative proxy sources are analyzed.

We demonstrated that the approach is highly accurate in most of the scenarios tested. Regardless of the availability of true or proxy admixture sources, our approach correctly assigned ancestral proportions, with a low associated error. Moreover, in the rare cases of error, ASAP appointed a group closely related (Fst % 0.01) to the source. Furthermore, ASAP can accurately assign ancestry proportions also in case of minor contributions as low as 10%. However, when only proxy sources are available it tends to overestimate the major component.

More importantly, ASAP performs well even when pseudo-haploid data with missing variants are analyzed, with a maximum assignment error of 3%. Specifically, when the admixture sources have a split time > 24 KYA, ASAP error estimates are lower than 0.014. On the other hand, for closely related admixture sources, the per-sample misassignment can reach 20% when multiple, closely related putative sources are available.

Compared to other global ancestry assignment tools, the approach is faster in terms of runtime while being as accurate (ChromoPainter) or more accurate than other tools (qpAdm) and, most importantly, provides ancestry estimates based on a straightforward formulation of user-defined ancestry sources with no need for in- or out-groups.

When tested on a real dataset of ancient and modern Eurasian genotypes, ASAP confirmed the trends in ancestry composition observed in previous research, providing relevant information on the complex scenario of the continent. Notably, it estimated significant gene flows after the Neolithic period in Aegean Bronze Age populations and confirmed previous findings about the shared origin of WHG ancestry in British and Iberian farmers.

Our approach relies on the assumption that the target group is indeed admixed. However, the target group might fall within a given cline for a demographic scenario different from admixture. This method should be used as an exploratory tool and subsequent analyses, such as formal tests for admixture, should be performed to test the admixture event further.

A future area of work is to explore and evaluate how the ASAP approach can also be applied to other summary statistics, such as IBS and/or Fst estimates, avoiding the evaluation of the number of PCs to be used.

## Materials and Methods

### DATASETS

#### Simulated dataset

##### • Genotype data with no missingness

We used a simulated genotype dataset from Molinaro et al. composed of 13 simulated demes with different population sizes and split times ranging from 250 to 4,000 generations, to represent a simplified scenario for current European (EUR 1-3), East Asian (ASN 1-3) and African (AFR 1-7) groups and 7 sister groups characterized by a split time from their closest population of 100 generations (Molinaro et al., 2021). The data simulation was carried out with mspms and following a modified Van Dorp et al. model (Kelleher et al., 2016; van Dorp et al., 2015). The initial dataset consisted of eight admixed groups, obtained by combining pairs of the simulated Ghost populations (GST), all with ancestry proportions of 70%-30%. The pairs of admixing GST were selected in order to cover a broad spectrum of split times. Specifically, we simulated admixture groups whose sources split time span from 75 KYA to 9 KYA, six sources shared a bottleneck event and for three of these, we simulated an additional one. The initial set also comprised one admixed group characterized by a three-way admixture with the proportions of 60%-20%-20%, with African-like, European-like, and Asian-like ancestries, respectively.

We simulated an additional three-way admixture group, using the same highly divergent sources as above, but different ancestry proportions, namely: 40% for the African-like ancestry, and 20% for both the European and Asian-like ancestries. To test models with strongly imbalanced ancestry proportions, we also simulated three groups with 90%-10% and three groups with 95%-05% ancestry proportions. In this case, as well, we chose the admixture sources (GST) to cover a broad spectrum of split times.

All admixture simulations were carried out with admix-simu (https://github.com/williamslab/admix-simu), creating 50 individuals per each population, using a constant recombination rate (1.25 x 10^-8^) and admixture time of 100 generations (Molinaro et al., 2021). We obtained a simulated dataset of 4,745,025 SNPs, 20 non-admixed, and 16 admixed groups. After filtering for minor allele frequency with PLINK (maf 0.01), the final dataset comprised 284.249 SNPs (Chang et al., 2015). PC analyses were performed on the final dataset, projecting the admixed target samples on the scaffold built from the non-admixed groups.

##### Ancient (Pseudo-haploid) data

To mimic the data quality of ancient DNA, we manipulated the simulated set by introducing both missing data as well as using pseudo-haploid genotypes. In each population, we introduced a variable missing rate (from 10% to 50%) in randomly selected positions, so every 10 individuals would be characterized by 10, 20, 30, 40 or 50% of missing data. Secondly, we created pseudo-haploid genomes by randomly selecting at each locus one allele and assigning it as a homozygous genotype, eventually obtaining for each simulated group 100 pseudo-haploid genomes from the original 50 diploid individuals. The missingness proportions were maintained after pruning.

PCA was performed after filtering for minor allele frequency (maf 0.01). For the pseudo-haploid datasets containing missing data, pruning was also performed (PLINK v1.9 indep-pairwise 50 10 0.1)(Chang et al., 2015).

After the filtering, the bim file of the modern simulated dataset contains 284.249 SNPs, the one in which only the real sources are pseudo-haploid has 135.211 SNPs, while the one in which the sources are also pseudo-haploid has roughly 100.000 SNPs.

##### Real modern and ancient dataset

We downloaded 1240K+HO dataset (version V52.2, https://reich.hms.harvard.edu/allen-ancient-dna-resource-aadr-downloadable-genotypes-present-day-and-ancient-dna-data) in EIGENSTRAT format. Such dataset includes present-day and ancient DNA data and converted it into PLINK format using convertf (Patterson et al., 2006).

Starting from the .anno file and following Aneli et al. method (Aneli et al., 2021), we created a list of ancient and modern samples to keep from the 1240K+HO dataset.

In particular, only the ancient ones (those that in the “Full date “column did not have the string “present”) coming from Western Eurasian countries (latitudes higher than 22 and longitudes between − 15 and 60) and the Mbuti individuals from Congo (string “Mbuti” in the “Genetic.ID” column) (Bergström et al., 2020; Patterson et al., 2012) were selected. Then we removed individuals that in the “Group.ID” column have the string “Ignore”. Finally, we kept only those whose “Assessment” column contained the string “PASS”. After removing any duplicates, we created a preliminary plink file with only 1240K+HO ancient samples to whom we added other ancient samples taken from other published datasets (Aneli et al., 2022; Lazaridis et al., 2022; Posth et al., 2021; Reitsema et al., 2022).

For the modern samples, we selected individuals coming from Western Eurasian countries with latitudes higher than 22 and longitudes between − 15 and 60, but removing those coming from Uzbekistan, Kazakhstan, Algeria, Morocco, Tunisia, Libya, as well as some populations from Russia and others showing “Ignore” within “Group.ID”. Then, we selected the samples flagged as “PASS” in the “Assessment” column. In this way, we created a preliminary plink file with only the 1240K+HO modern individuals to which we merged other modern-day samples taken from the Raveane et al. dataset (Raveane et al., 2019).

From this modern dataset, we extracted only the autosome chromosome SNPs and excluded those monomorphic (--maf .00001) and with more than 5% missing data (--geno .05) using PLINK (Chang et al., 2015).

Then we extracted the bulk of variants built on our modern dataset from the ancient dataset and finally merged all with PLINK1.9 excluding the ancient samples with less than 20,000 SNPs (N=1,381). To assess the relatedness, we computed the kinship coefficient pi-hat, expressing the probability that two randomly selected alleles, belonging to the same locus, are identical-by-descent (IBD) for any two samples. Using --genome --min 0.35 in PLINK1.9 we selected pairs of samples showing a pi-hat of at least 0.35, retaining 7,312 individuals. We then filtered out samples with duplicated IDs retaining the one with a higher number of SNPs. We carried out a Principal Component Analysis (PCA) using smartpca, through which we filtered out outliers obtaining a dataset of 6,408 samples. Our final real dataset, containing 4,740 ancient and 1,668 modern samples with 206,363 SNPs, was converted to the EIGENSTRAT format using convertf.

### Principal Component Analysis (PCA)

PCA was performed using smartpca (version 16000) available in the EIGENSOFT package (Patterson et al., 2012).

The admixed (or target) populations were always projected, regardless of the dataset used. In the case of datasets with pseudo-haploid or ancient individuals, we projected all samples’ genotypes onto the principal components inferred from the diploid/modern individuals using the lsqproject: YES option. For each analysis on the simulated and real genotype datasets, we run 10 PCs. Furthermore, we performed the same analysis using different numbers of PCs, in order to assess the performance of ASAP (see Supplementary Text 1)

### Non Negative Least Squares (NNLS)

To perform the Non-Negative Least Squares, we used the NNLS function, as described in (Hellenthal et al., 2014; Leslie et al., 2015; Ongaro et al., 2019), which is an adaptation of the Lawson–Hanson NNLS implementation of Non-Negative Least Squares (Lawson and Hanson, 1995) available in the statistical software package R 3.5.1 (R Core Team, 2020).

We applied NNLS both population-wise and individual-wise. To estimate the NNLS population-wise, we estimated the average of each of the population PCs and then applied NNLS on the resulting vector. On the other hand, to estimate the NNLS individual-wise, the PCs of each individual were maintained as separate vectors. Error values reported in the text were calculated as the absolute average difference between the observed and the expected proportion assignment. We used a block jackknife approach to resampling our set and estimate the standard errors. Given that the simulated set consisted of only chromosome 1, we could not use chromosomes as blocks, as usually it’s done when the entire genome is available. We thus estimated the number of SNPs available after filtering and divided them into 20 blocks. For each resampling step, we removed one of such blocks and performed PCA on the remaining ones.

Standard errors were estimated on chromosome-based jackknife replications (Büsing et al., 2011).

### qpAdm

To validate the results obtained from ASAP, we performed the most widely used approach to assess the ancestry components and the relative proportions of the admixed population: qpAdm programs in the ADMIXTOOLS package (Patterson et al., 2012).

For each admixed individual, we tested as “left” populations all the possible true sources and used all the others as right populations. Subsequently, we selected the inference characterized by the largest p-value, irrespectively to their significance. Although there are many ways to harness qpAdm to obtain more reliable results, we decided to use a strategy comparable to the other tools harnessed here.

### Rye

We applied Rye (Conley et al., 2023) converting the PCA output obtained by smartpca using a custom R script. We performed five different rounds using the first ten PCs.

### ChromoPainter

ChromoPainter (CP) (Lawson et al., 2012) was applied using the unlinked (-u) model, where, for each SNP in the target, we assign a score of 1/K to each reference haplotype that carries the same allele, where K is the total number of reference haplotypes that carry the same allele.

### Analysis of Eurasian ancient and modern genotype data

We carried out a PCA computing 10 Principal Components (PCs) per each individual in our final dataset projecting ancient samples on the top of present-day genome variability (N = 1668). We used smartpca version 16000 with autoshrink lsqproject options for this analysis. Subsequently, we selected ancient individuals previously analyzed in Lazaridis et al., 2022 (Lazaridis et al., 2022) and filtered out samples with less than 180K SNPs. This resulted in a dataset comprising 1,380 ancient donors, of which 30 were chosen as recipients for ASAP. The selection of recipient samples was based on the five main ancestral sources identified in Lazaridis 2022 (Lazaridis et al., 2022), namely Western Hunter-Gatherers (WHG), Eastern Hunter-Gatherers (EHG), Caucasus Hunter-Gatherers (CHG), Anatolian Neolithic, and Levant Neolithic. ASAP was run using 10 PCs, and the estimates were correlated with F4admix results using all samples combined, as well as stratified by different ancestral sources. Pearson correlation analysis was performed using the ggpubr library in R. To explore ancestry over time, we visualized single ancestry trends by either selecting individuals as indicated in the publication or by visually inspecting populations present in (Lazaridis et al., 2022) figures.

## Acknowledgements

LDG and MV were supported by #NEXTGENERATIONEU (NGEU) and funded by the Ministry of University and Research (MUR), National Recovery and Resilience Plan (NRRP), project MNESYS (PE0000006) – A Multiscale integrated approach to the study of the nervous system in health and disease (DN. 1553 11.10.2022). FM was supported by Fondazione con il Sud (2018-PDR-01136) and by the Italian Ministry of University and Research (2022P2ZESR). MV was supported by the Italian Ministry of University and Research (2022E8NN2N). FS is a PhD student within the European School of Molecular Medicine (SEMM). GH was supported by the Wellcome Trust (224575/Z/21/Z). LP is funded by the Italian Ministry of University and Research (PRIN 2022B27XYM). LM and TK were supported by KU Leuven BOF-C24 grant ZKD6488 C24M/19/075 and FWO grant G0A4521N (TK).

We would like to thank Nicole Soranzo for advice on the final stage of the manuscript preparation.

