## Supplementary figures and images for "Fast and reliable ancestral reconstruction on ancient genotype data with non-negative Least square and Principal Component Analysis"

### Supplementary Figure 1

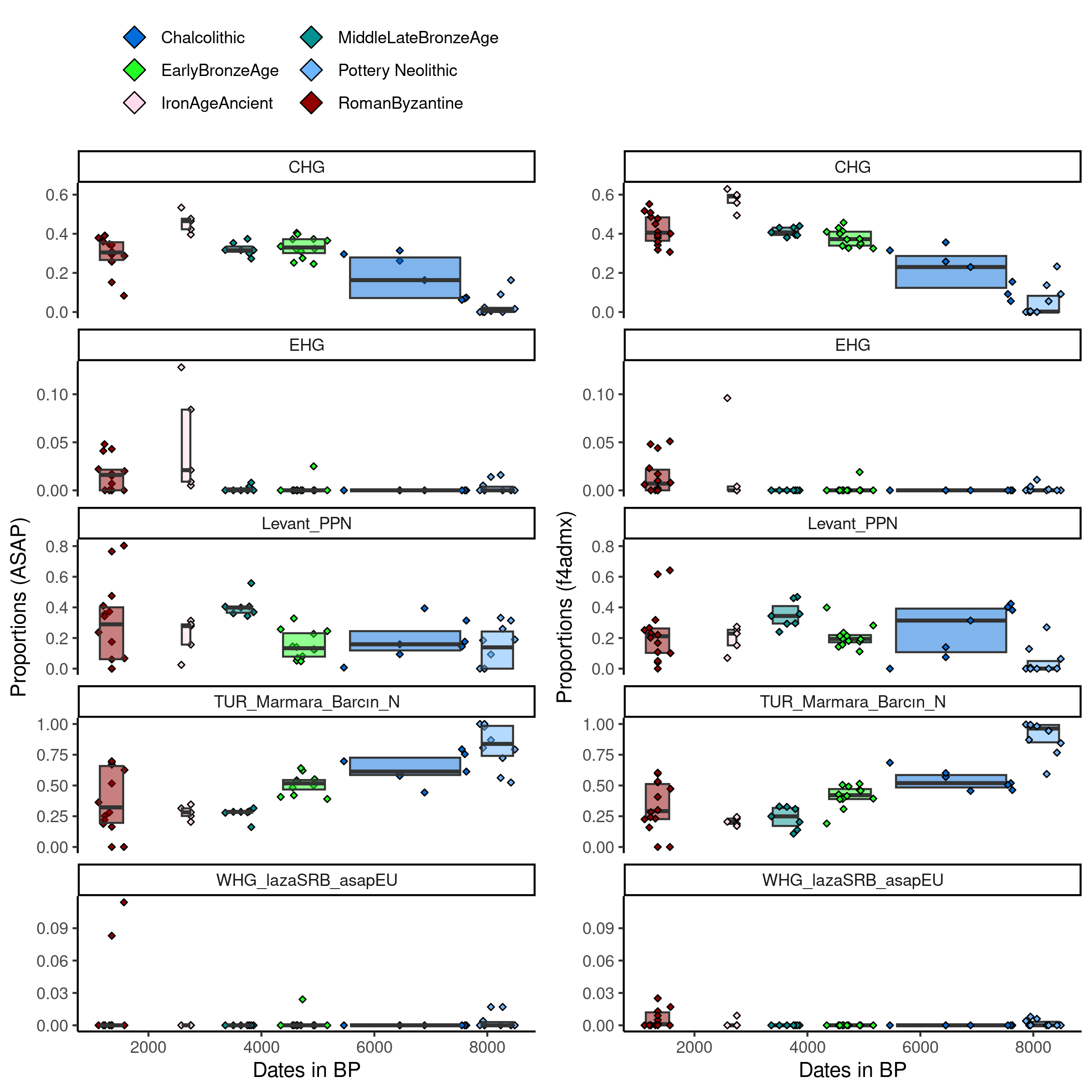

### Supplementary Figure 2

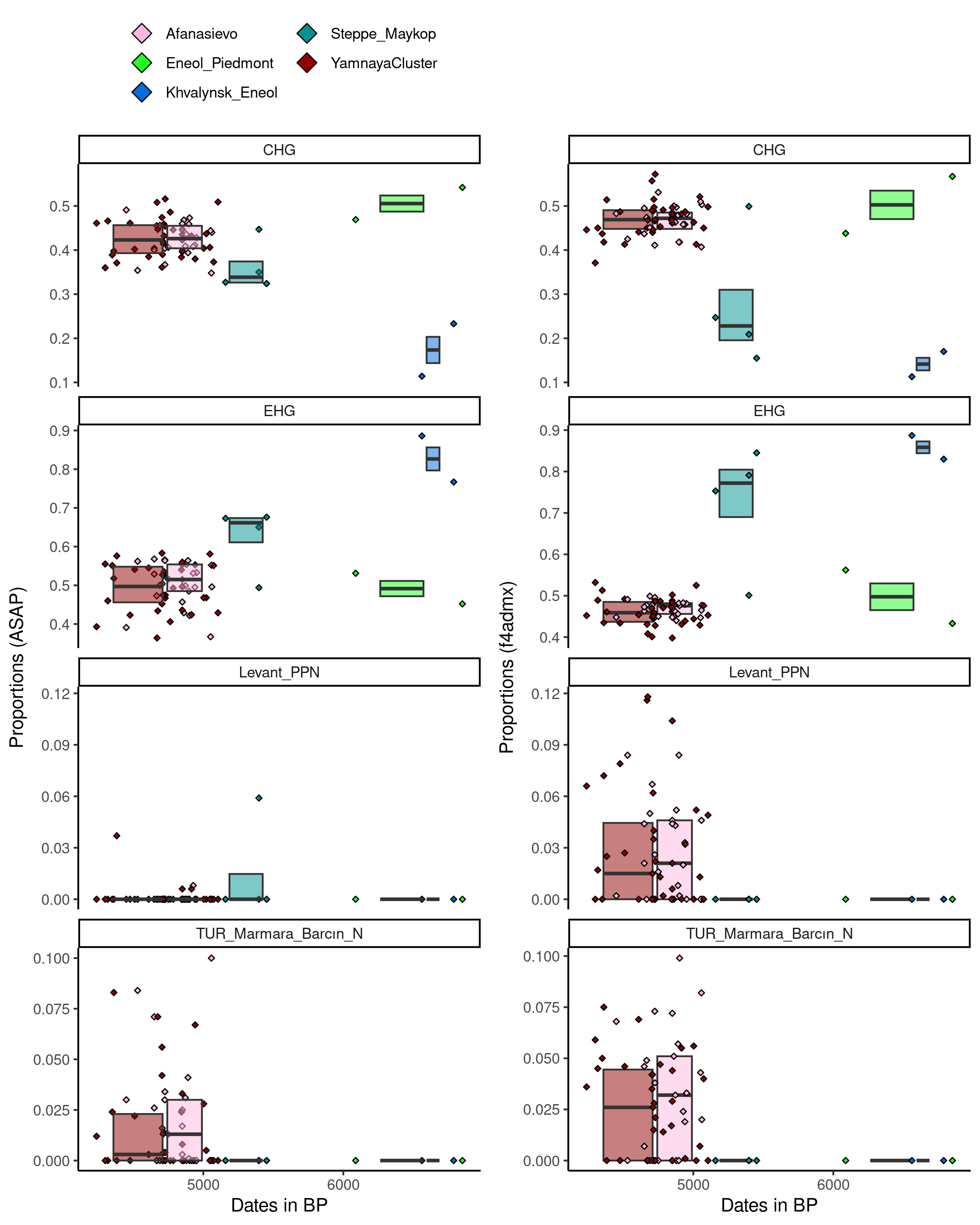

### Supplementary Figure 3

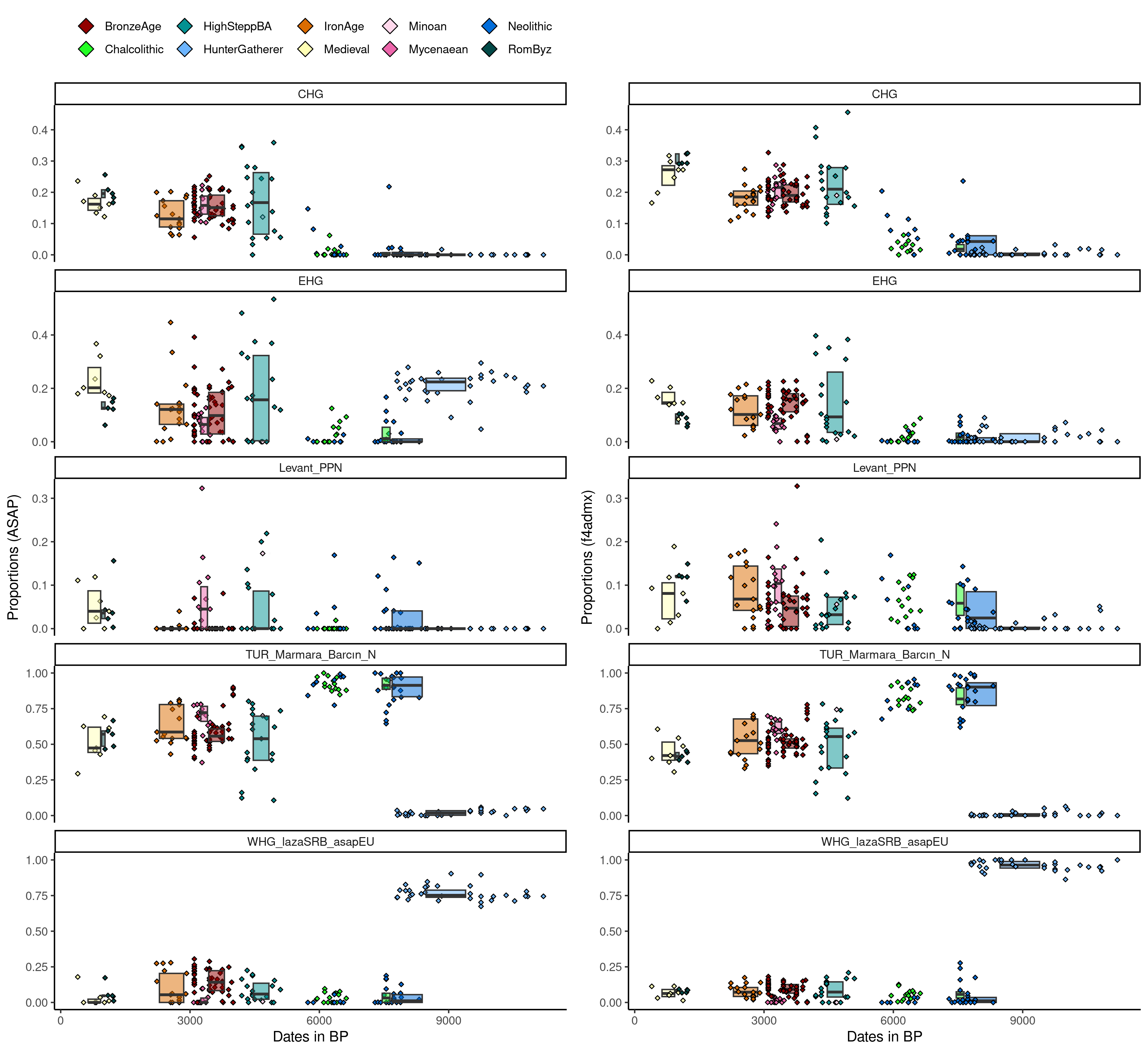

### Supplementary Figure 4

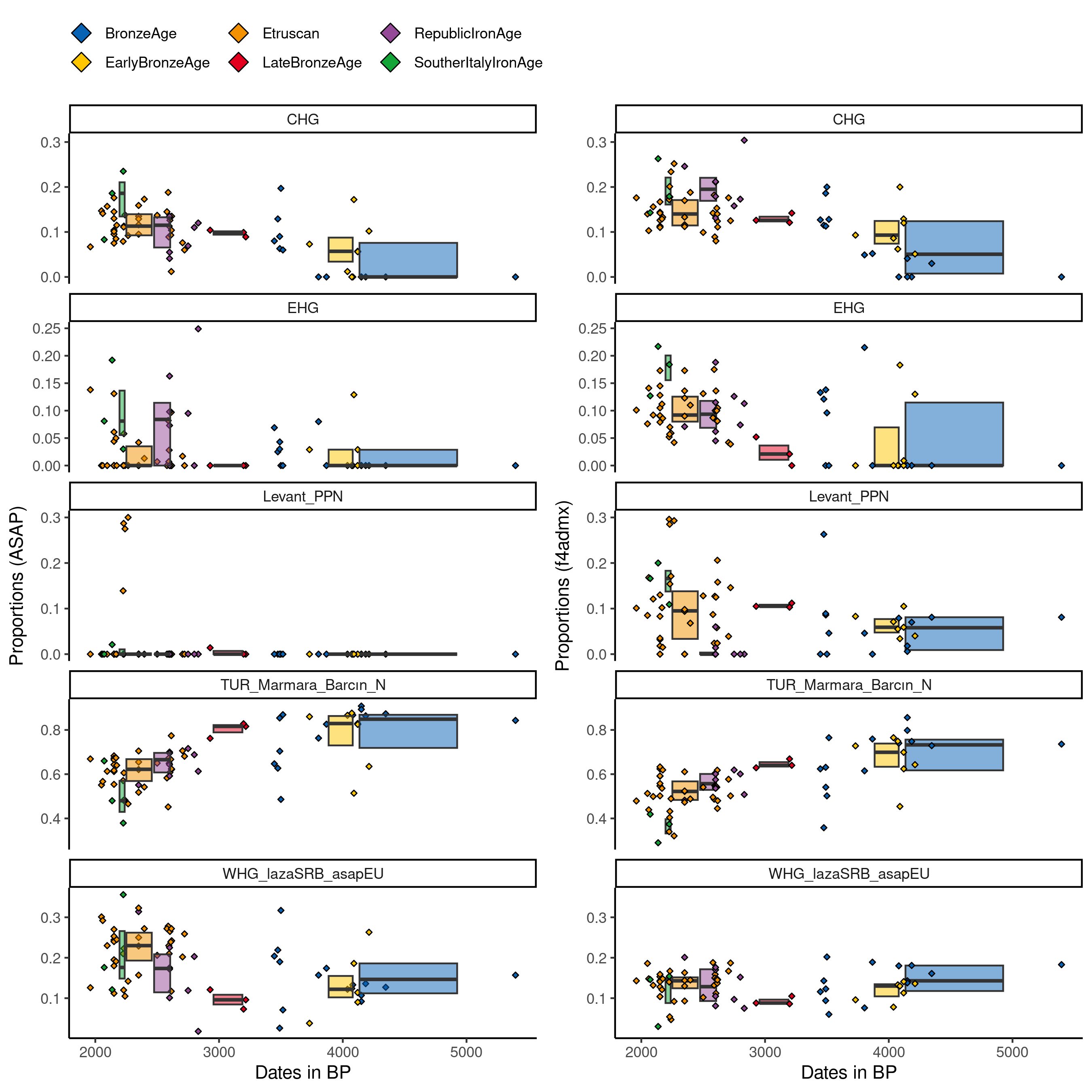
